# Improving estimates of negative selection in human genome using CAPS

**DOI:** 10.1101/2024.01.23.576817

**Authors:** Mikhail Gudkov, Loïc Thibaut, Eleni Giannoulatou

## Abstract

Despite ongoing efforts, variant interpretation in disease sequencing studies is often hindered by the lack of well-established ways of determining the potential pathogenicity of genetic variation, especially for understudied classes of single-nucleotide variants (SNVs). Population genetics methods offer an attractive solution to this problem by enabling the assessment of the effects of SNVs through their distributions in human populations.

For instance, negative selection is known to shift site-frequency spectra of genetic variation, thus affecting the ratio of singleton variants. It has been shown that the extent of negative selection can serve as a proxy for deleteriousness. An example of this approach is the Mutability-Adjusted Proportion of Singletons (MAPS) metric. Although MAPS proves a useful instrument for the assessment of selection-based deleteriousness in SNVs, it is highly sensitive to the calibration of the singletons-by-mutability model, which results in potentially biased estimates for some classes of variants.

Building up on the methodology used in MAPS, we developed a novel metric of negative selection in the human genome — CAPS, or Context-Adjusted Proportion of Singletons. Compared to its predecessor, CAPS provides estimates of negative selection that are less biased and have more accurate confidence intervals. CAPS inherits some of the same features that make MAPS useful for studying SNVs, yet the key difference of our method is the complete elimination of the mutability layer in the model, which makes the metric more robust and reliable.

We believe that CAPS holds promise for improving the discovery of new disease-variant associations in clinical and research settings.

## Introduction

Most disease-variant association studies these days are hindered by the lack of well-established ways of variant prioritisation, which remains one of the key challenges in modern genomic analyses. The problem of variant prioritisation can be reduced to comparing variants based on their potential deleteriousness. However, for many classes of genetic variation their overall deleteriousness remains poorly quantified.

With the ever-increasing size of open databases of human genetic variation, population genetics methods have the potential to provide the means of variant prioritisation that are based on patterns of genetic variation that occur naturally in human populations. One of the key considerations when analysing variants through the population genetics approach is the variability in mutation rates, which is known to affect variant analysis (Karczewski & Martin, 2020). Indeed, as some variants have higher mutability (the ability to spread easily in populations) than others, variants with high mutability are less likely to be rare and can reach saturation in large genomic databases. This creates a need for the use of the finite sites model and mutability correction (Harpak et al., 2016).

Transversions, CpG transitions and non-CpG transitions constitute the 3 main types of single-nucleotide variants. Transversions (for example, CGG to CGT) have much lower mutation rates than CpG transitions, because the mutations that they create are more complex biochemically. It has been shown that trinucleotide sequence context is sufficient to account for a large proportion of variability in mutation rates and that the baseline mutability level can be obtained from the mutation rates of trinucleotide sequences in noncoding regions (Harpak et al., 2016; Samocha et al., 2014). Using this approach, each variant falls into one of 104 possible groups, as there are 32 trinucleotide contexts with each having 3 possible mutations of the middle nucleotide, with the additional 8 groups coming from the 4 CpG contexts’ medium and high levels of methylation.

A particular example of a population genetics method that utilises this approach is the Mutability-Adjusted Proportion of Singletons (MAPS) metric (Karczewski et al., 2020; Lek et al., 2016), which can be used as a tool for estimating negative selection and deleteriousness. The general assumption in MAPS is that if a particular variant is damaging, it will be rare in the population, because purifying selection will be trying to remove it. MAPS scores are calculated as the scaled excess or deficit of singletons (variants with allele count of 1), where the expected number is derived based on context sequence from a singletons-by-mutability model calibrated on a relatively neutral class of variants. Even though it has been shown that intergenic variants can be used for calibration (Findlay et al., 2022), synonymous variants have traditionally been preferred for this purpose. Importantly, given then the complexity of mutability estimation, there is no straightforward way of modelling mutation rates; however, attempts have been made to incorporate additional mutability-related annotations in the singletons-by-mutability model (Findlay et al., 2022).

Here we show that the mutability layer in MAPS makes its estimates biased, with the severity of the bias varying depending on the variant type composition in the dataset of interest. We present CAPS, the Context-Adjusted Proportion of Singletons metric, as a drop-in replacement for MAPS with better performance and unbiased estimates of negative selection.

## Results

We developed CAPS, a novel metric of negative selection where the expected level of rare variation is derived on a per-context basis instead of using a mutability-based model.

### Improved estimates of negative selection

As shown in Figures 1 and S1, the estimates of MAPS carry a strong bias coming from the singletons-by-mutability model, which is an integral part of MAPS’ design. Specifically, in MAPS, transversion variants are more likely to be assigned a missense-level negative selection score merely due to their low mutation rates. It is important to note, however, that this bias may not be noticeable when MAPS is calculated over large variant sets. CAPS eliminates this mutability bias completely, as its design does not include the mutability correction layer (Figure 2). As a result, unlike MAPS, CAPS can be used safely to study a wide range of different classes of genetic variation, regardless of the variant type composition in the variant set of interest.

**Figure 1.**
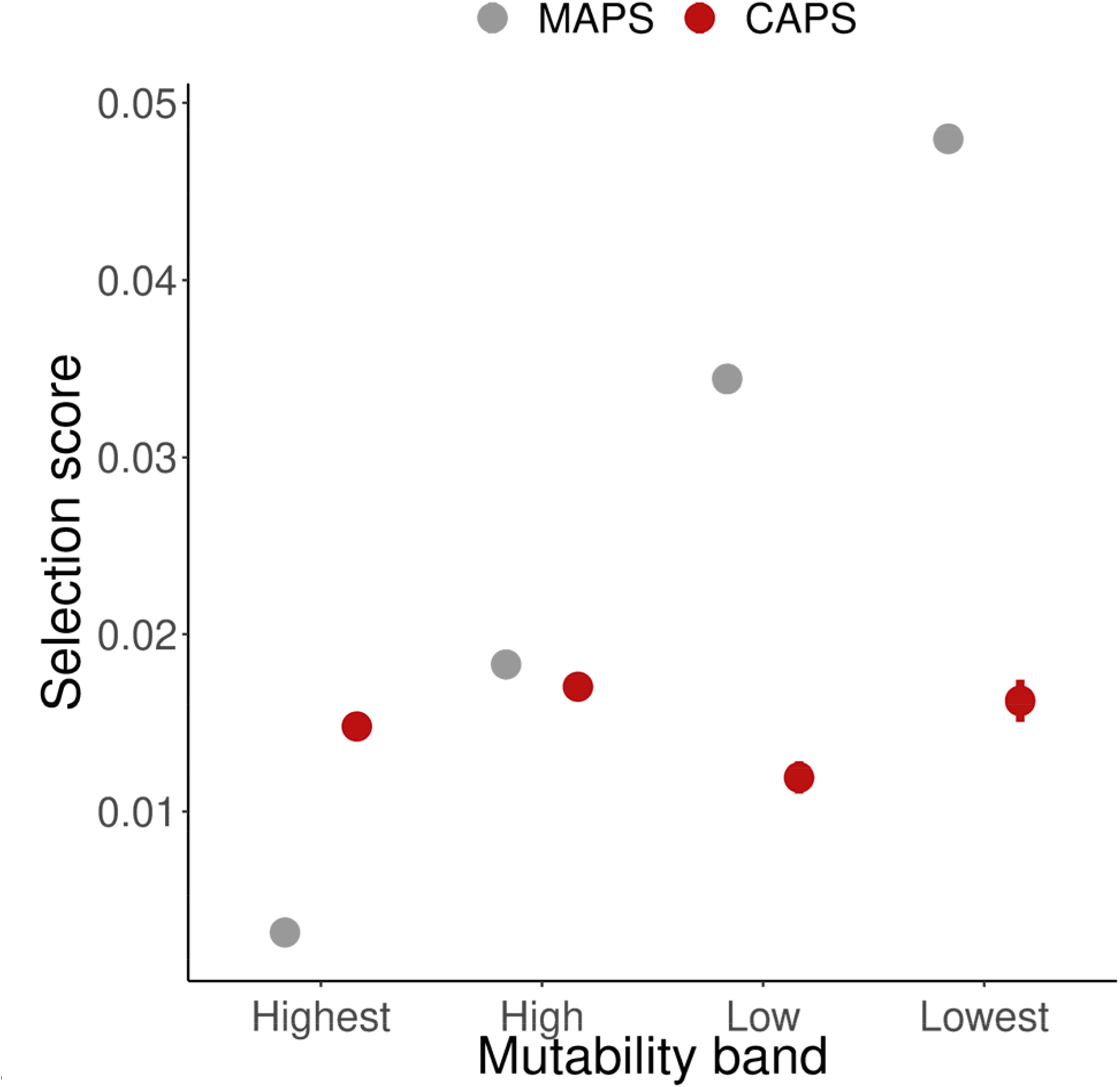
Estimates of negative selection in CAPS and MAPS by mutability band: “Lowest” (0-25%), “Low” (25-50%), “High” (50-75%), “Highest” (75-100%). All QC-compliant WES variants.

**Figure 2.**
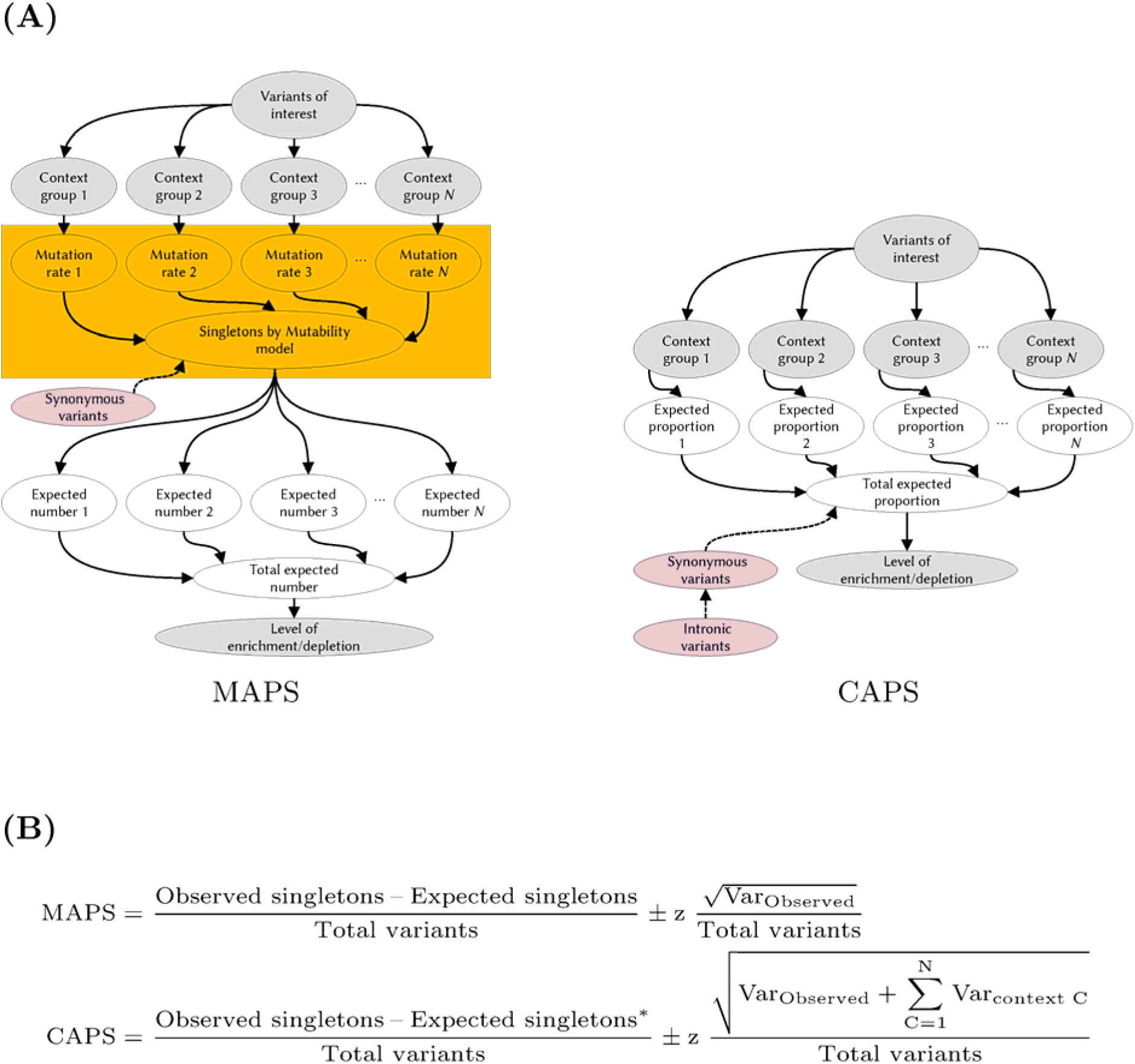
Differences between CAPS and MAPS in the derivation of the scores. In CAPS, the complexity of the model is reduced via complete elimination of the mutability layer. Star (*) indicates that in CAPS the expected level of rare variation is derived in a different way compared to MAPS.

As shown in Figure 3, CAPS’ scores of negative selection are highly consistent with those of MAPS, with the estimates of the two metric agreeing for all categories of variants when either the exome or genome frequency data from the gnomAD database are used. Specifically, both MAPS and CAPS capture the same upward trend in the deleteriousness of synonymous, missense and predicted loss-of-function (pLoF) variants. The major difference in the results observed is in the confidence intervals of the estimates, which are more accurate in CAPS (Table S1).

**Figure 3.**
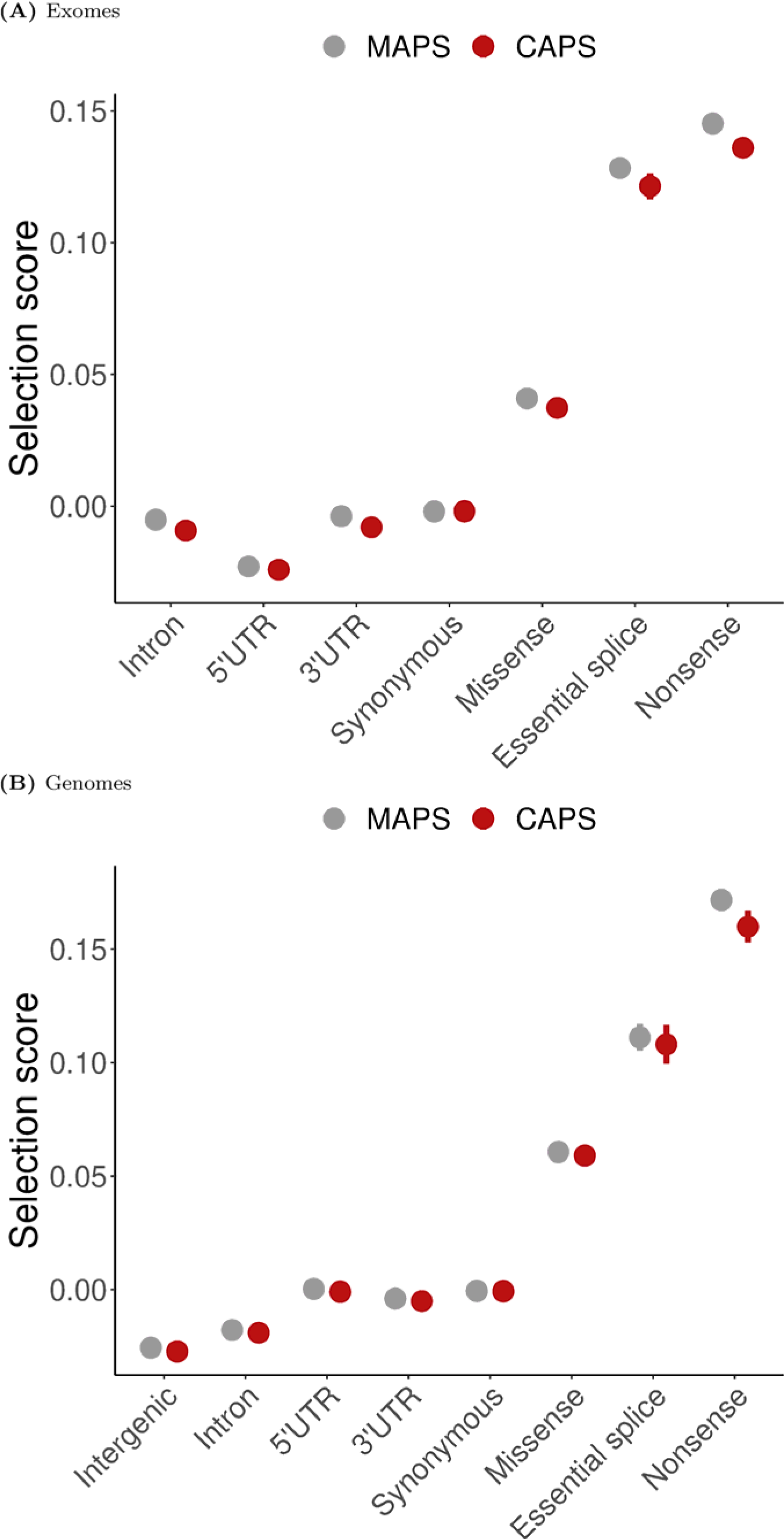
Original (MAPS) and corrected (CAPS) estimates of negative selection in SNVs by variant class for exomes (A) and genomes (B). All QC-compliant variants.

### CAPS as a drop-in replacement for MAPS

To demonstrate the applicability of CAPS we sought to apply it to those classes of variants which had been previously studied using MAPS. Figures 4 and 5 show how CAPS can be used as a drop-in replacement for MAPS, using reproduced scores from previously reported analyses in upstream open reading frames (uORFs) and near-splice regions, respectively (Blakes et al., 2022; Lord et al., 2019; Whiffin et al., 2020). As evident from the figures, the CAPS’ negative selection scores show good concordance with those estimated using MAPS. These results confirm that uORF uAUG-creating and some uORF stop-removing variants are subject to strong negative selection and that this selection is dependent on the effects that these variants induce and contextual information. Our results also confirm that variants affecting intron-exon junctions are particularly deleterious.

**Figure 4.**
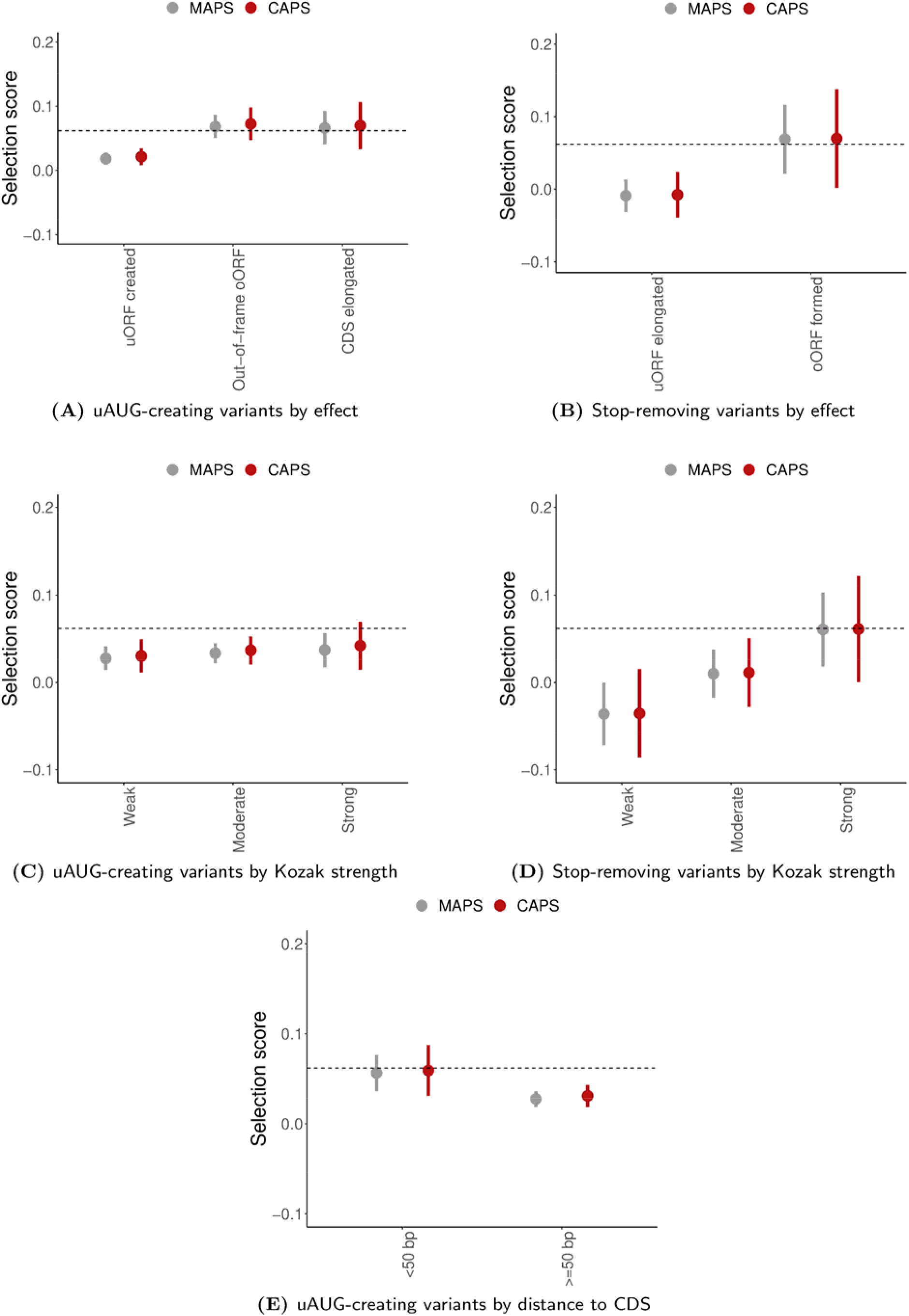
Corrected estimates of negative selection in uAUG-creating and stop-removing variants in uORFs.

**Figure 5.**
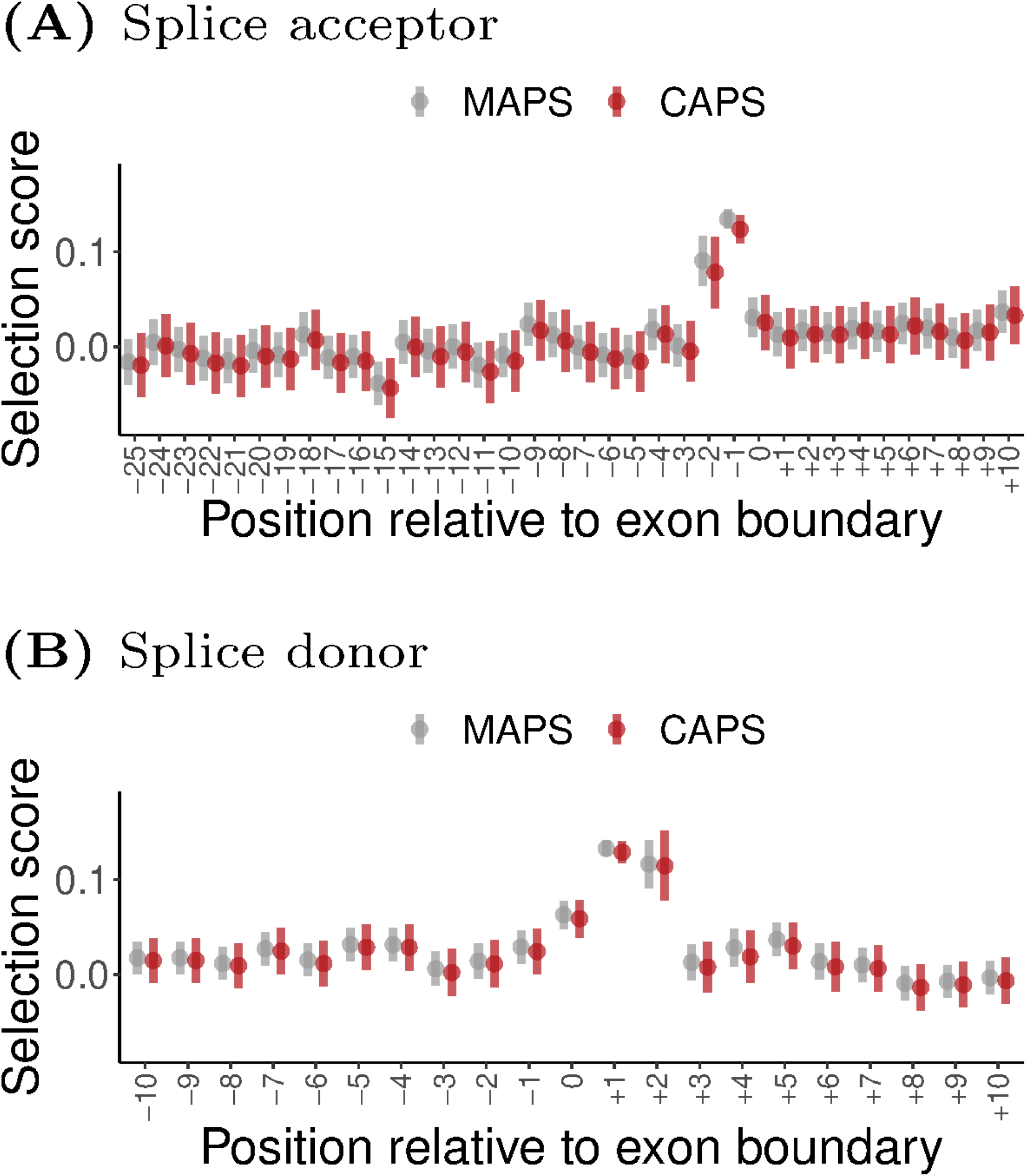
Corrected estimates of negative selection in variants affecting near-splice positions.

### CAPS captures background selection in constrained genes

To check the validity of CAPS’ corrected estimates of selection, we compared CAPS and MAPS scores in sets of variants with increasing level of gene constraint using LOEUF, a metric of intolerance to variation based on the deficit of pLoF variants in a gene (Karczewski et al., 2020). To minimise the effect of targeted selection, we limited our dataset to synonymous variants and stratified all variants by gene intolerance level.

Figure 6 shows that both metrics are sensitive enough to capture the background selection that affects synonymous variants in constrained genes and have striking similarity in the scores. Wider confidence intervals can be seen for CAPS.

**Figure 6.**
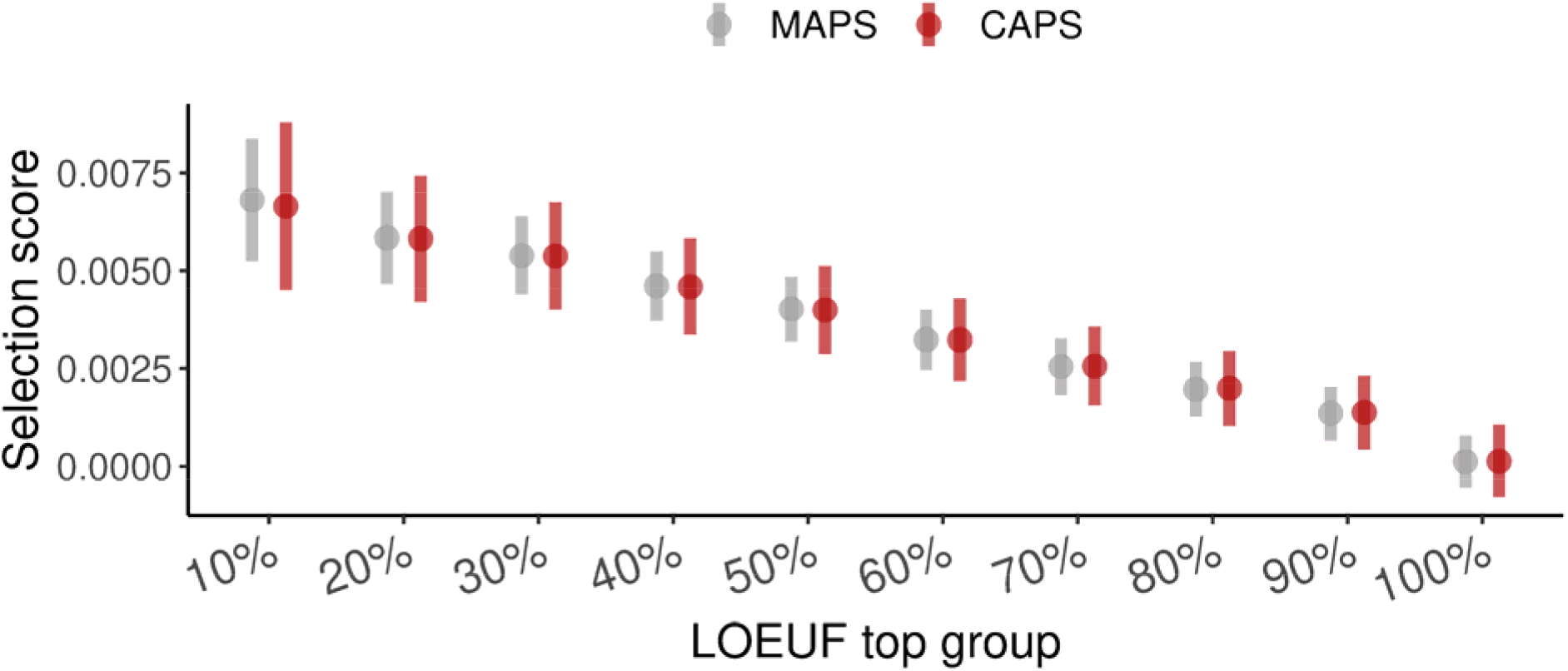
CAPS and MAPS scores in genes with different levels of LOEUF-defined constraint. Synonymous QC-compliant WES variants only. “10%”: top 10% most constrained genes; “100%”: all genes.

## Discussion

In this work, we introduced CAPS, a novel metric of selection-based deleteriousness and a drop- in replacement for MAPS. CAPS eliminates the mutability bias that was present in its predecessor and performs reliably even on small variant sets. Besides, CAPS’ confidence intervals are calculated in a more accurate way compared to the simplified intervals used in MAPS.

Overall, we believe that our proposed context-based approach is superior compared with using mutability as a proxy for the estimation of the expected level of rare variation, as mutation rates modelling proves a difficult task (Findlay et al., 2022). Even though it is possible to reduce bias of MAPS towards transversion variants by modifying the fit of the model, we argue that this approach can be seen as ad hoc and not future-proof, considering that every new release of gnomAD would require refitting the model.

We believe that CAPS holds promise to reveal new insights into understudied classes of variation with unknown or poorly quantified deleteriousness, especially for smaller classes of variants. Compared with complex, machine learning-based pathogenicity predictors, the design of CAPS is simpler and more interpretable. We believe that this transparency will help researchers to better understand the potential impact of genetic variants on human health.

## Materials and methods

### Score calculation procedure

CAPS is a new metric of negative selection with the expected level of rare variation derived on a per-context basis from synonymous variants.

Even though, as mentioned earlier, it has been shown that intergenic variants may be used as an alternative class of variants for the estimation of the baseline level of rare variation (Findlay et al., 2022), we had insufficient evidence that intergenic variants could be used interchangeably with synonymous variants for the purpose of model calibration without any significant effect on the results.

Given that in synonymous variants 4 out of 104 contexts are never observed, due to how the genetic code works, we were initially unable to use synonymous variants to calculate CAPS scores for some classes of genetic variation where those contexts were observable. We, therefore, approximated the expected proportions of singletons for those missing contexts from intronic variants using a probit regression model. This enabled us to apply CAPS to all classes of variants, including those with all 104 contexts present. Importantly, CAPS can be calibrated for both genomes and exomes, just like MAPS.

One of the key advantages of using per-context estimates of the expected proportion of singletons over the singletons-by-mutability approach is that each context can be seen as a binomial random variable, which allows the calculation of per-context variance in a mathematically sound way. This, in turn, results in wider and more realistic confidence intervals compared with MAPS’ binomial confidence intervals for the mean with the assumption of normality. Figure 2B explains this idea further through the comparison of the mathematical formulas for the calculation of CAPS and MAPS scores.

CAPS scores can be calculated using either a simple method, which is based on the total variance in the observed number of singletons, or using the posterior predictive distribution (PDD) of CAPS. The actual values of the scores are identical between the two methods; however, the produced confidence intervals differ: the intervals estimated using the PDD method are wider, as they take into the additional uncertainty around the probabilities that are used to calculate the expected level of singletons for each context (see Table S1).

### Data

For model calibration and all analyses we used 125,748 exomes (WES) and 15,708 genomes (WGS) from gnomAD v2.1.1, filtered using the same QC criteria as in the original 2020 gnomAD flagship paper (Karczewski et al., 2020). The total number of QC-compliant variants per class is shown in Tables S2 and S3, with additional per-context statistics shown in Tables S4 and S5.

## Web resources

- gnomAD: https://gnomad.broadinstitute.org

## Data and code availability

The code generated during this study is available at https://github.com/VCCRI/CAPS

## Supplemental data

Supplemental data include 1 figure and 5 tables.

## Acknowledgements

We thank Professor Daniel MacArthur, Director of the Centre for Population Genomics, for his valuable feedback on this work.

## References

Blakes, A. J. M., Wai, H. A., Davies, I., Moledina, H. E., Ruiz, A., Thomas, T., Bunyan, D., Thomas, N. S., Burren, C. P., Greenhalgh, L., Lees, M., Pichini, A., Smithson, S. F., Taylor Tavares, A. L., O’Donovan, P., Douglas, A. G. L., Whiffin, N., Baralle, D., Lord, J., … Group, D. W. (2022). A systematic analysis of splicing variants identifies new diagnoses in the 100,000 genomes project. Genome Medicine, 14.

Findlay, S. D., Romo, L., & Burge, C. B. (2022). Quantifying negative selection in human 3’ UTRs uncovers constrained targets of RNA-binding proteins. bioRxiv.

Harpak, A., Bhaskar, A., & Pritchard, J. K. (2016). Mutation rate variation is a primary determinant of the distribution of allele frequencies in humans. PLOS Genetics, 12(12), 1–22.

Karczewski, K. J., Francioli, L. C., Tiao, G., Cummings, B. B., Alföldi, J., Wang, Q., Collins, R. L., Laricchia, K. M., Ganna, A., Birnbaum, D. P., Gauthier, L. D., Brand, H., Solomonson, M., Watts, N. A., Rhodes, D., Singer-Berk, M., England, E. M., Seaby, E. G., Kosmicki, J. A., … MacArthur, D. G. (2020). The mutational constraint spectrum quantified from variation in 141,456 humans. Nature, 581(7809), 434–443.

Karczewski, K. J., & Martin, A. R. (2020). Analytic and translational genetics. Annual Review of Biomedical Data Science, 3(1), 217–241.

Lek, M., Karczewski, K. J., Minikel, E. V., Samocha, K. E., Banks, E., Fennell, T., O’Donnell-Luria, A. H., Ware, J. S., Hill, A. J., Cummings, B. B., Tukiainen, T., Birnbaum, D. P., Kosmicki, J. A., Duncan, L. E., Estrada, K., Zhao, F., Zou, J., Pierce-Hoffman, E., Berghout, J., … Williams, A. L. (2016). Analysis of protein-coding genetic variation in 60,706 humans. Nature, 536(7616), 285–291.

Lord, J., Gallone, G., Short, P. J., McRae, J. F., Ironfield, H., Wynn, E. H., Gerety, S. S., He, L., Kerr, B., Johnson, D. S., McCann, E., Kinning, E., Flinter, F., Temple, I. K., Clayton-Smith, J., McEntagart, M., Lynch, S. A., Joss, S., Douzgou, S., … Deciphering Developmental Disorders study, on behalf of the. (2019). Pathogenicity and selective constraint on variation near splice sites. Genome Research, 29(2), 159–170.

Samocha, K. E., Robinson, E. B., Sanders, S. J., Stevens, C., Sabo, A., McGrath, L. M., Kosmicki, J. A., Rehnström, K., Mallick, S., Kirby, A., Wall, D. P., MacArthur, D. G., Gabriel, S. B., DePristo, M., Purcell, S. M., Palotie, A., Boerwinkle, E., Buxbaum, J. D., Cook, E. H., … Daly, M. J. (2014). A framework for the interpretation of de novo mutation in human disease. Nature Genetics, 46(9), 944–950.

Whiffin, N., Karczewski, K. J., Zhang, X., Chothani, S., Smith, M. J., Evans, D. G., Roberts, A. M., Quaife, N. M., Schafer, S., Rackham, O., Alföldi, J., O’Donnell-Luria, A. H., Francioli, L. C., Armean, I. M., Banks, E., Bergelson, L., Cibulskis, K., Collins, R. L., Connolly, K. M., … Ware, J. S. (2020). Characterising the loss-of-function impact of 5’ untranslated region variants in 15,708 individuals. Nature Communications, 11(1), 1–12.

