## Supplementary Material for "Improving estimates of negative selection in human genome using CAPS"

### Supplemental Figures

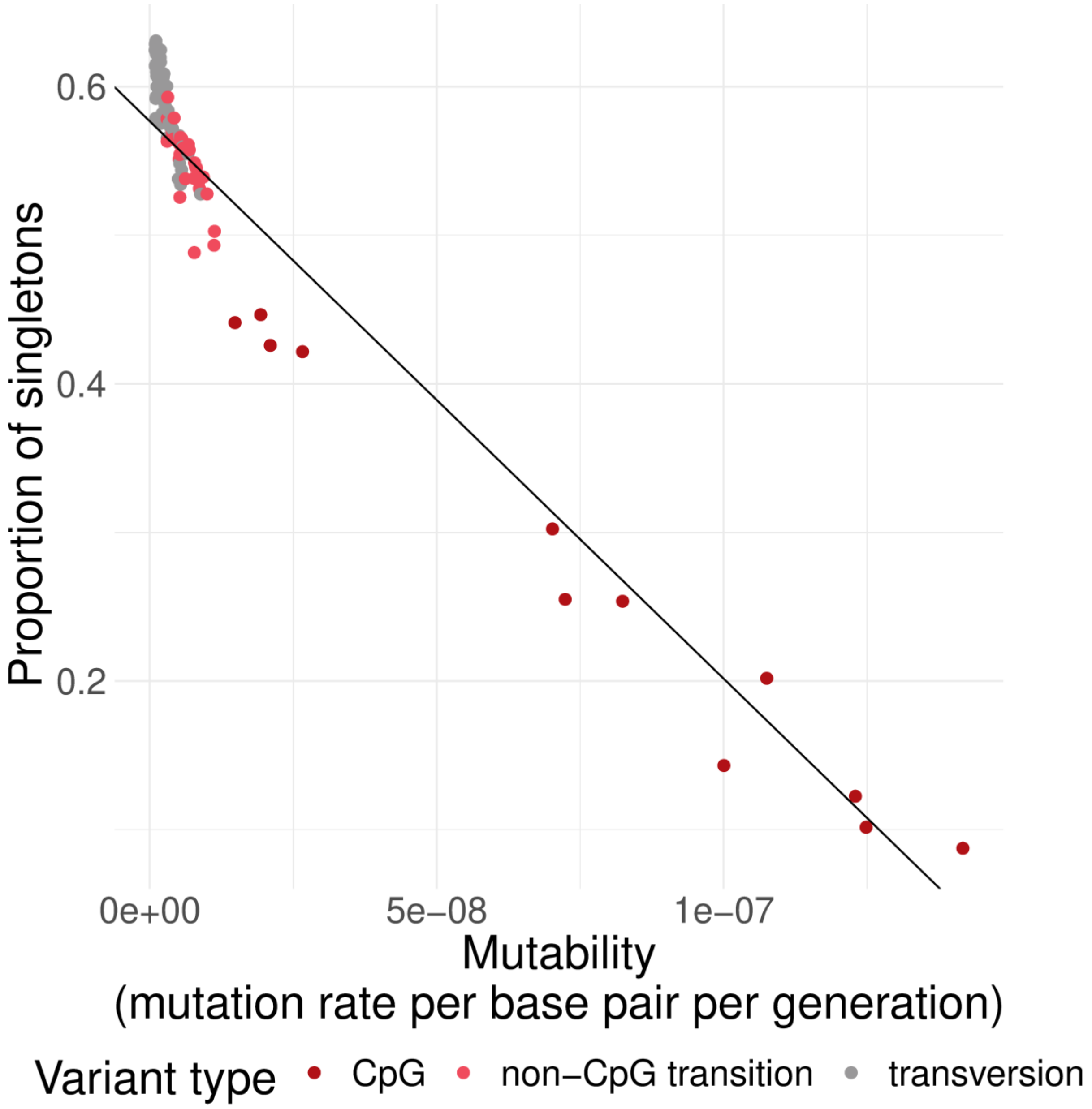


Figure S1. MAPS' singletons-by-mutability model calibrated on synonymous variants. The MAPS model underestimates the real proportion of singletons for transversions, while overestimating it for CpGs, though this may not be critical when averaging over large numbers of variants.

### Supplemental Tables

Table S1. Increase in the size of confidence intervals in CAPS and CAPS-PDD compared to MAPS by variant class. All QC-compliant WES variants.

| **Subset** | **Model** | **CI increase** |
|---|---|---|
| Intron | CAPS | 39% |
| Intron | CAPS-PDD | 136% |
| 5’UTR | CAPS | 40% |
| 5’UTR | CAPS-PDD | 77% |
| 3’UTR | CAPS | 38% |
| 3’UTR | CAPS-PDD | 79% |
| Synonymous | CAPS | 37% |
| Synonymous | CAPS-PDD | 100% |
| Missense | CAPS | 38% |
| Missense | CAPS-PDD | 179% |
| Essential splice | CAPS | 44% |
| Essential splice | CAPS-PDD | 84% |
| Nonsense | CAPS | 38% |
| Nonsense | CAPS-PDD | 80% |

Table S2. The total number of variation observed in each variant class for exomes.

| **Class** | **Total** |
|---|---|
| Intron | 2436006 |
| 5’UTR | 108405 |
| 3’UTR | 160419 |
| Synonymous | 2173989 |
| Missense | 4547687 |
| Essential splice | 65907 |
| Nonsense | 132974 |

Table S3. The total number of variation observed in each variant class for genomes.

| **Class** | **Total** |
|---|---|
| Intergenic | 48346752 |
| Intron | 63771613 |
| 5’UTR | 267647 |
| 3’UTR | 1384125 |
| Synonymous | 627102 |
| Missense | 1155661 |
| Essential splice | 19646 |
| Nonsense | 28977 |

Table S4. Per-class context statistics for exomes.

| **Class** | **Contexts observed** | **Transversion contexts** | **Weighted mean mutability, $10^{-8}$** |
|---|---|---|---|
| Intron | 104 | 0.615 | 1.5 |
| 5’UTR | 104 | 0.615 | 1.3 |
| 3’UTR | 104 | 0.615 | 1.8 |
| Synonymous | 100 | 0.6 | 2.6 |
| Missense | 104 | 0.615 | 2.3 |
| Essential splice | 90 | 0.6 | 1.0 |
| Nonsense | 69 | 0.71 | 3.0 |

Table S5. Per-class context statistics for genomes.

| **Class** | **Contexts observed** | **Transversion contexts** | **Weighted mean mutability, $10^{-8}$** |
|---|---|---|---|
| Intergenic | 104 | 0.615 | 1.8 |
| Intron | 104 | 0.615 | 2.1 |
| 5’UTR | 104 | 0.615 | 1.4 |
| 3’UTR | 104 | 0.615 | 2.3 |
| Synonymous | 100 | 0.6 | 4.1 |
| Missense | 104 | 0.615 | 3.5 |
| Essential splice | 100 | 0.62 | 1.3 |

| Class | Contexts observed | Transversion contexts | Weighted mean mutability, $10^{-8}$ |
|---|---|---|---|
| Nonsense | 64 | 0.688 | 4.1 |
